# Streamlining the isolation of fungal hyphae: A semi-automated approach for soil substrates

**DOI:** 10.1101/2025.03.14.643305

**Authors:** Isabelle Elisabeth Metzen, Marcel Bucher

## Abstract

Extracting fungal hyphae with their natural associated microbiota from soil samples presents a significant challenge due to their small size, typically in the micrometer range, and the formation of dynamic fungal networks. Previous methods, which involved supplementing soil substrates with sand or glass beads, have proven technically challenging for large-scale field applications and tend to create artificial conditions that disrupt the plant-microbe-soil continuum. In this study, we introduce a semi-automated approach for efficiently extracting fungal hyphae from a variety of soil types, including natural loamy soils. The Sieving and Sucrose Centrifugation (SSC) technique enables the enrichment of fungal hyphae, spores, and their surface-associated bacteria, with subsequent analysis of hyphal length density and the identification of tightly attached surface bacteria via next-generation sequencing (NGS). A comparison of different hyphal extraction techniques revealed that the SSC method yielded maximal hyphal length density. Furthermore, the SSC approach effectively enriched fungal hyphae from a highly diverse community, establishing a dependable method for advancing soil microbial research.

## 1 Introduction

Plant tissues and their immediate surroundings host a diverse array of microorganisms, including bacteria, protists, viruses, oomycetes and fungi. Most fungi in natural soils form hyphae - filamentous structures that extend and branch throughout the soil. These hyphae play a crucial role in nutrient acquisition, facilitating the uptake and transport of essential elements. They also contribute to the spread of fungal networks, support spore formation, and enable the colonization of plant roots, thereby shaping soil microbial communities and plant-fungal interactions. Some fungi, such as yeasts, either completely lack hyphae or have them in a highly reduced form, living primarily as single cells (Singara Charya, 2015). These non-hyphal fungi make up only a small fraction of the total fungal biomass in soil, whereas hyphal-forming fungi dominate and play a crucial role in decomposing organic matter and driving nutrient cycling (Treseder & Lennon, 2015; van der Heijden et al., 2015; Hagenbo et al., 2022). Additionally, some fungi, such as *Mucor* species, exhibit dimorphism, switching between yeast-like and hyphal forms in response to environmental conditions. This ability enhances their adaptability, allowing them to colonize diverse habitats and interact more effectively with hosts (Boyce & Andrianopoulos, 2015). Among filamentous fungi which colonize plant roots, arbuscular mycorrhizal fungi (AMF) of the subphylum *Glomeromycotina* are well-studied, while fungi of the subphylum *Mucoromycotina* have been more recently explored (Orchard et al., 2017a). Fungi from both taxa form mutualistic symbioses with their plant host and extend their extraradical hyphae into the soil, accessing nutrient patches beyond the plant’s root system. They absorb nutrients such as phosphorus (P) in the form of inorganic orthophosphate (Pi) or nitrogen (N) and deliver them to plant roots in exchange for carbon in the form of sugars and lipids (Pfeffer et al., 1999; Trépanier et al., 2005; Jiang et al., 2017; Keymer et al., 2017; Prout et al., 2024; Spatafora et al., 2016). The full diversity of fungi and associated microbes capable of directly providing nutrients for plant uptake remains incompletely understood.

AMF depend on interactions with soil bacteria for Pi uptake due to their limited ability to mineralize soil organic P (Tisserant et al., 2013; Zhang et al., 2014; Wang et al., 2017). AMF extraradical hyphae (ERH) extend into the soil, guiding bacteria to nutrient-rich patches, and enhance nutrient acquisition. These hyphae secrete carbon-rich compounds, such as fructose, to attract and support specific bacteria, modulating their metabolic activity (Zhang et al., 2016; Zhang et al., 2022). For example, AMF have been shown to increase bacterial phosphatase activity, which improves plant P nutrition (Zhang et al., 2018; Zhang et al., 2020). Mutual alteration of functional traits between AMF and soil-borne bacterial strains, which colonize the ERH, leads to a long-term and balanced cross-kingdom interaction (Anckaert et al., 2024).

Although only a subset of soil bacteria attach to or enter fungal hyphae (Toljander et al., 2007; Scheublin et al., 2010), microbial analyses often lack consistent distinctions between the hyphal compartments: the hyphosphere (soil influenced by ERH), the hyphal endosphere (inside hyphae), and the hyphoplane (hyphal surface). Bacterial communities at the AMF hyphoplane differ significantly from those in the hyphosphere and bulk soil (Gahan et al., 2015; Zhang et al., 2018b; Wang et al., 2019; Zhou et al., 2020; Emmett et al., 2021; Wang et al., 2022). However, it remains unclear how far the hyphal sphere of influence extends into the surrounding soil and differentiates microbial communities. Early work in 1991 suggested that fungal hyphae can alter pH and P levels in their vicinity (Li et al., 1991). This raised the question of whether hyphal effects extend far enough to differentiate the microbial community in the hyphosphere, not just in the hyphoplane, from the bulk soil. Insights into this question emerged from a few studies, highlighting that the hyphosphere varies in fungal and bacterial community composition compared to bulk soil (Nuccio et al., 2013; Xu et al., 2018; Qin et al., 2022; Emmett et al., 2021). Thus, hyphal presence has a detectable impact on microbial composition in the surrounding soil volume, appropriately termed the hyphosphere. A study investigating all three compartments - hyphoplane, hyphosphere, and bulk soil - further confirmed that the bacterial community in the hyphosphere differs in the abundance of *Burkholderiaceae*, *Sandaracinaceae*, *Chloroflexi*, and *Acidibacter* from that in the bulk soil (Emmett et al., 2021).

Cross-kingdom research, particularly focusing on the interactions between AMF hyphae and bacteria or other fungi, is an emerging field of research that has in the past received relatively little attention, likely due to the inherent difficulty of isolating hyphae, that typically range in size from 2 to 20 µm, from natural substrates. So far, only a limited number of studies have explored whether roots and hyphae assemble distinct bacterial and fungal communities, despite strong indications that such differences exist. For instance, Zhang et al. (2024) reported that roots and AMF hyphae host distinct bacterial communities. However, the underlying mechanisms determining the extent to which plant genotypic variation influences these microbial communities remain largely unknown. One possible explanation could be differences in hyphal secretory activities, which may shape microbial assemblages in unique ways.

In field, greenhouse, and *in vitro* experiments aimed at characterizing microbial communities assembled by fungal hyphae, a compartmented system with a small-pore, root-impermeable nylon mesh is commonly used to separate roots from hyphae (Jakobsen and Rosendahl, 1990). Historically, methods for extracting fungal hyphae from soil began in the late 1940ies, when Jones et al. (1948) mixed soil samples into molten agar, spread them thinly on slides, and stained them for microscopic analysis. Later, various approaches, such as manual hyphal picking under the stereomicroscope, sieving, blending, or centrifugation, were developed or modified. Hanssen et al. (1974) introduced a filter-based method where blended soil was filtered, washed, and stained for hyphal quantification. Abbott et al. (1984) refined this technique, and Jakobsen et al. (1992) further improved it for sandy soils. Bingle and Paul (1986) developed a method combining wet filtration with sucrose density centrifugation, suitable for AMF hyphae and spore extraction. This method shows parallels to the methodology of Allen (1979), which was designed to isolate spores from soil. Awad and Pena (2023) recently adapted this method for hyphae from diverse soil types.

After hyphal extraction, hyphal length density can be manually measured using the visual gridline intersection (VGI) method (Newman, 1966; Tennant, 1975; Shen et al., 2016) or semi-automated tools like the Fiji plug-in AnaMorf (Barry et al., 2009; Schindelin et al., 2012) or the HyLength software (Cardini et al., 2020), respectively.

Several methodologies for hyphal extraction have been described, particularly in the 20th century, yet many studies have incorporated sand and glass beads into the soil of the root-free compartment to simplify the process (e.g., Jakobsen et al., 1992; Wang et al., 2019; Zhou et al., 2020; Emmett et al., 2021; Zhang et al., 2024). However, adding extra components to soil substrates is not a practical solution for large-scale field applications. Such modifications often create artificial conditions that can disrupt the natural interactions between plants, microbes, and soil, potentially altering the delicate balance of the plant-microbe-soil continuum.

Here, we introduce a semi-automated sieving and sucrose centrifugation (SSC) method for efficiently extracting fungal hyphae with attached microbes directly from various natural soils, including clay-rich field soils. This high-throughput method is based on Bingle and Paul (1986) and Jakobsen et al. (1992) and incorporates automated wet sieving with high shaking amplitude to remove loamy soil, followed by sucrose centrifugation for concentrating hyphae and spores. The SSC method enables hyphal length quantification and identification of microbes closely associated with fungal hyphae.

## 2 Material and Methods

Previously developed methods for the efficient extraction of fungal mycelium from various soil substrates were optimized and integrated. Using a sieving machine, the process was semi-automated to increase the number of samples processed per day while reducing the required manpower. Additionally, the goal was not only to optimize a method for hyphal length density estimation but also to develop a technique that preserves surface-attached microbes on fungal hyphae. This approach enables the subsequent identification of bacteria on the hyphoplane through NGS.

### Experimental set-up

Soil containing fungal hyphae was harvested from three field experiments conducted in 2019, 2020, and 2021 at the University of Cologne Field Station, located at coordinates 50.925617° N, 6.936641° E. Field plots were allocated for NK (-P), PK (-N), or NPK (full) fertilized silty or sandy loam. To spatially separate hyphae-associated compartments (hyphosphere, hyphoplane, and hyphal endosphere) from root-associated compartments (rhizosphere, rhizoplane, and root endosphere), hyphal containers (HCs) were used. These cylindrical plastic containers (Thermo Fisher Scientific, USA; 10077900 X12 jar, 125 mL PP, straight-walled, autoclavable) had a cut-out screw cap sealed with parafilm. They were inserted into the soil with a 21 μm nylon mesh facing the growing plant root, allowing fungal hyphae to extend into the container while preventing root intrusion. This setup ensures that microbial communities associated with fungal hyphae can be studied independently of direct root influence. Three HCs per plant were inserted at two depths (topsoil: 3–10 cm and subsoil: 23–30 cm) and two distances (2 cm and 9 cm) from the root system center (Figure 1A). The HCs were collected during the vegetative stage of *Zea mays*. To obtain the hyphosphere sample, the natural soil inside the hyphal container was collected. The hyphoplane sample was isolated using the SSC method, while bulk soil was sampled from an unplanted area of the field.

**Figure 1:**
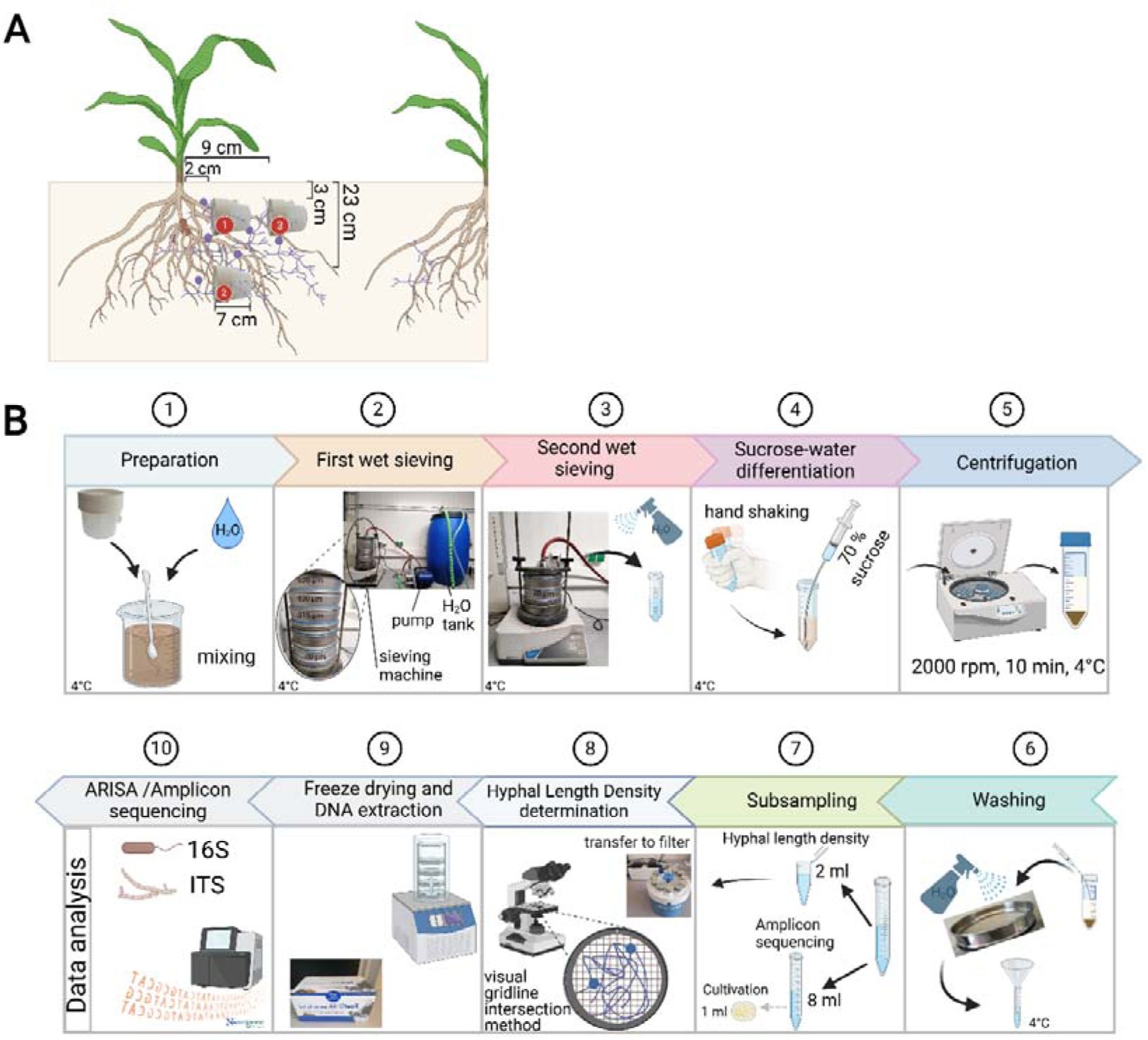
(A) Schematic illustrating the positioning of hyphal containers containing natural soil, fungal spores and mycelium and microorganisms, but excluding roots, in the fields. Each 7 cm-long plastic container creates individual hyphae-associated compartments. The containers are placed horizontally at two distances from the center of the root system and at two different depths in the soil for each plant. The figure is not to scale. (B) Stepwise procedure for isolation of hyphae from natural field soil. Hyphal containers were harvested and processed semi-automatically using the SSC method at 4°C. The method includes steps 1-10. Illustrations made with BioRender.com.

For practical reasons, an alternative substrate was used to develop the SSC method. This substrate, referred to as QSF, consisted of a quartz/soil mixture that had been previously planted with *Allium schoenoprasum* and inoculated with *Rhizophagus irregularis* spores (DAOM 197198). As a result, it contained a high density of fungal mycelium, making it well-suited for method optimization.

### Extraction of fungal hyphae

A combination of semi-automated wet sieving steps and sucrose centrifugation, termed the SSC method, was used to extract fungal hyphae from soil substrates directly or from soil within hyphal containers (Figure 1B). Soil samples were mixed with sterile, 4°C-cold water to detach fungal hyphae from dense soil particles. The samples were placed on the first sieve of the sieving convoy. 4°C-cold water was pumped through a low-pressure water pump into the wet sieving machine, which consisted of a water-blast pipe and the sieving convoy. The water pressure and amplitude were set to 2.5 bar and 75%, respectively. The samples underwent an initial sieving process lasting 8 minutes. The first three sieves with the widest mesh sizes were removed, and the remaining material was sieved for an additional 6 minutes. The particles retained on the 40 μm sieve were collected and transferred into 50 mL microcentrifuge tubes, with the samples kept on ice. The volume of each microcentrifuge tube was adjusted to 20 mL by adding H_₂_O as needed. The solutions were hand-shaken to stir up the particles, and equal amounts of 70% sucrose were injected beneath the aqueous solution. The samples were then centrifuged (2000 rpm for 10 minutes at 4°C). Hyphae and spores were collected from the interface of the two solutions. They were placed on a 40 μm sieve, gently washed with water, and subsequently collected in a centrifuge tube. The effectiveness of UV light treatment, hypochlorite treatment, and water rinsing in preventing DNA contamination on sieves for subsequent sample processing was evaluated. PCR-based detection using the 515F (GTGCCAGCMGCCGCGGTAA) and 806R (GGACTACHVGGGTWTCTAAT) primer pair, which targets the V4 region of the 16S rRNA gene for bacterial and archaeal community analysis, was used. Amplification of the 16S and ITS2 regions revealed that water rinsing was the most effective method for minimizing bacterial and fungal contamination on sieves. In agarose gel electrophoresis, no amplification was visible in the DNA extracted from the water that passed through the rinsed sieve. Therefore, these water samples were not used for amplicon sequencing.

### Estimation of hyphal length density

To measure hyphal length density (HLD) (fungal hyphae in cm per hyphal compartment), 2 mL of the solution containing hyphae and spores was filtered onto a nitrocellulose membrane (pore size: 1.2 μm) using a manifold holding system connected to a vacuum pump. A vacuum was applied to remove excess liquid from the filter. The retained hyphae and spores were then stained by immersing the filter in an ink solution for 20 minutes. After staining, the ink was aspirated through the vacuum, and the sample was rinsed with ddH_₂_O. Once dried, the intersections of hyphae with the gridlines were counted at 13 sites per sample under a Leica DM 1000 LED microscope at 10× magnification. Hyphal length density was subsequently calculated following the methods of Tennant (1975) and Shen et al. (2016).

### Comparison of methods for hyphal length counting

We compared two semi-automated methods for HLD counting (Barry et al., 2009; Cardini et al., 2020) with the manually performed VGI method (Supplementary Figure S14) (Tennant, 1975; Shen et al., 2016). The first semi-automated approach was performed with the Fiji plug-in AnaMorf (Barry et al., 2009; Barry et al., 2015; Schindelin et al., 2012) following the publishers’ recommendations while the second used the software *HyLength* (Cardini et al., 2020). To evaluate these methods, fungal hyphae and spores were extracted using the SSC method from different sample sizes (100 g, 50 g and 25 g) of the QSF. This substrate, rooted by *Allium schoenoprasum*, provided a sufficient amount of AMF hyphae for analysis.

### DNA extraction and microbiome sequencing

The remaining water solution containing hyphae was lyophilized. DNA was extracted from the dried hyphae and spores using the Fast DNA Spin Kit for Soil (MP Biomedicals, Eschwege, Germany) according to the manufacturer’s protocol, with the modification that homogenization was performed twice. ITS2 amplicons for fungi were generated using the ITS3 (GCATCGATGAAGAACGCAGC) and ITS4 (TCCTCCGCTTATTGATATGC) primer pair, while 16S V4 amplicons were obtained using the 515F (GTGCCAGCMGCCGCGGTAA) and 806R (GGACTACHVGGGTWTCTAAT) primer pair (White et al., 1990; Caporaso et al., 2011). PCR amplification, sample quality control, and library generation were performed by the sequencing provider Novogene (Munich, Germany). In general, six replicates per condition and controls (H_₂_O, empty kit control) were sequenced on the NovaSeq PE250 instrument, with a sequencing depth of 50,000 raw reads per sample. The sequencing reads were demultiplexed, and primers and barcodes removed by the sequencing provider.

The raw sequences were processed with the DADA2 pipeline v1.8.0 and the phyloseq package in RStudio, to obtain amplicon sequencing variants (ASVs) (Callahan et al., 2016; McMurdie and Holmes, 2013).

### Scanning electron microscopy

To verify bacterial attachment to hyphae after applying the SSC method, samples were analyzed using a scanning electron microscope (SEM; FEI Quanta 250 FEG) at 20 kV. To further confirm bacterial presence on AM fungal hyphae, two additional approaches were employed: (1) PCR-based detection using the 515F (GTGCCAGCMGCCGCGGTAA) and 806R (GGACTACHVGGGTWTCTAAT) primer pair, and (2) DAPI (4′,6-diamidino-2-phenylindole) staining.

### Comparison to the modified aqueous blending and sieving method (BSM) and the differential water/sucrose centrifugation method (WSCM)

The hyphal yield obtained via the SSC method was compared to that from a modified aqueous blending and sieving method (Jakobsen et al., 1992) and a modified differential water/sucrose centrifugation method (Allen et al., 1979; Bingle and Paul, 1986). Hyphae and spores were extracted from 80 g of a quartz/soil mixture following respective protocols, and hyphal length density (HLD) was measured using the VGI method.

For the sucrose centrifugation method, soil from hyphal containers was mixed with 400 ml ddH_₂_O, passed through a 315 μm sieve, and collected on a 40 μm sieve. The aqueous sample was underlayered with 70% sucrose, centrifuged at 2000 rpm for 10 min. Hyphae and spores were collected from the sucrose-water boundary layer, washed, and transferred to a 15 ml tube for further analysis.

For the blending and sieving method, the quartz/soil mixture was blended with 250 ml of ddH_₂_O for 30 seconds, briefly shaken, and allowed to settle for 1 min. A 2 ml supernatant sample was then taken for HLD measurement.

### General statistics and figure generation

Statistical analyses were conducted in RStudio (version 1.4.1). Data distribution was assessed using the Shapiro-Wilk test of normality (Shapiro.test function) (Royston, 1982, 1995). Depending on the distribution, statistical tests were selected, including ANOVA followed by Tukey’s HSD test, the Wilcoxon test, or the Kruskal-Wallis test (Bauer, 1972; Hollander & Douglas, 1973; de Mendiburu & de Mendiburu, 2019). The agricolae package in R was used for ANOVA with Tukey’s test (p < 0.05) and the Wilcoxon test (p < 0.05, FDR corrected). Figures were generated using the ggplot function from the ggplot2 package (Wickham, 2016) for data visualization. Correlograms were produced using the ggcor function in ggplot2, based on Spearman correlation. Venn diagrams were generated by using the ggvenn_pq function from the MiscMetabar package (Taudière, 2015). Figures were further refined and finalized in Inkscape (v.0.294) or BioRender for improved presentation. Schematic illustrations were designed using BioRender to ensure clarity and consistency.

## 3 Results and Discussion

We developed the SSC method with two primary objectives: (1) to efficiently extract fungal hyphae and (2) to investigate the microbial communities attached to them. The method was specifically designed to preserve the integrity of both spores and hyphae as much as possible, ensuring minimal disruption during extraction.

In the initial step, optimal parameters, such as water pressure and sieving amplitude, were determined using the QSF mixture, which contained a high abundance of fungal hyphae (see Materials and Methods). An amplitude above 75% caused damage to spores, while a water pressure of 2.5 bar was sufficient to extract hyphae while minimizing water consumption (Supplementary Figure S1A). Based on these findings, the parameters were set to 2.5 bar water pressure and 75% amplitude. The amount of remaining soil organic matter and residual material (including sand, roots, and other debris) varied depending on soil type, with sandy soils leaving more residual material than soils with higher clay content (Supplementary Figure S2).

Since field soil often contains stones with diameters of up to 15 mm - classified as medium gravel (USDA Soil Taxonomy size classification, C. P, 1977) - and was not pre-filtered before being added to the hyphal containers in our experiment, we tested the mechanical impact of medium gravel on the integrity of hyphae and spores. To do so, medium gravel was incorporated into a stone-free QSF substrate containing sand (Supplementary Figure S1B). Despite the presence of gravel, both hyphae and the majority of fungal spores remained intact.

A comparison of the SSC method with modified versions of the previously described aqueous blending and sieving method (BSM) and the differential water/sucrose centrifugation method (WSCM) showed that the SSC method yielded higher amounts of extraradical hyphae from field soil than the other two methods (Figure 2). Under our conditions, the manual VGI method provided the most accurate hyphal counts and demonstrated effective scalability across different sample sizes. In contrast, semi-automated counting using the Fiji plug-in AnaMorf (Barry et al., 2009; Barry et al., 2015; Schindelin et al., 2012) or the HyLength software (Cardini et al., 2020) showed lower consistency (Supplementary Figure S3).

**Figure 2:**
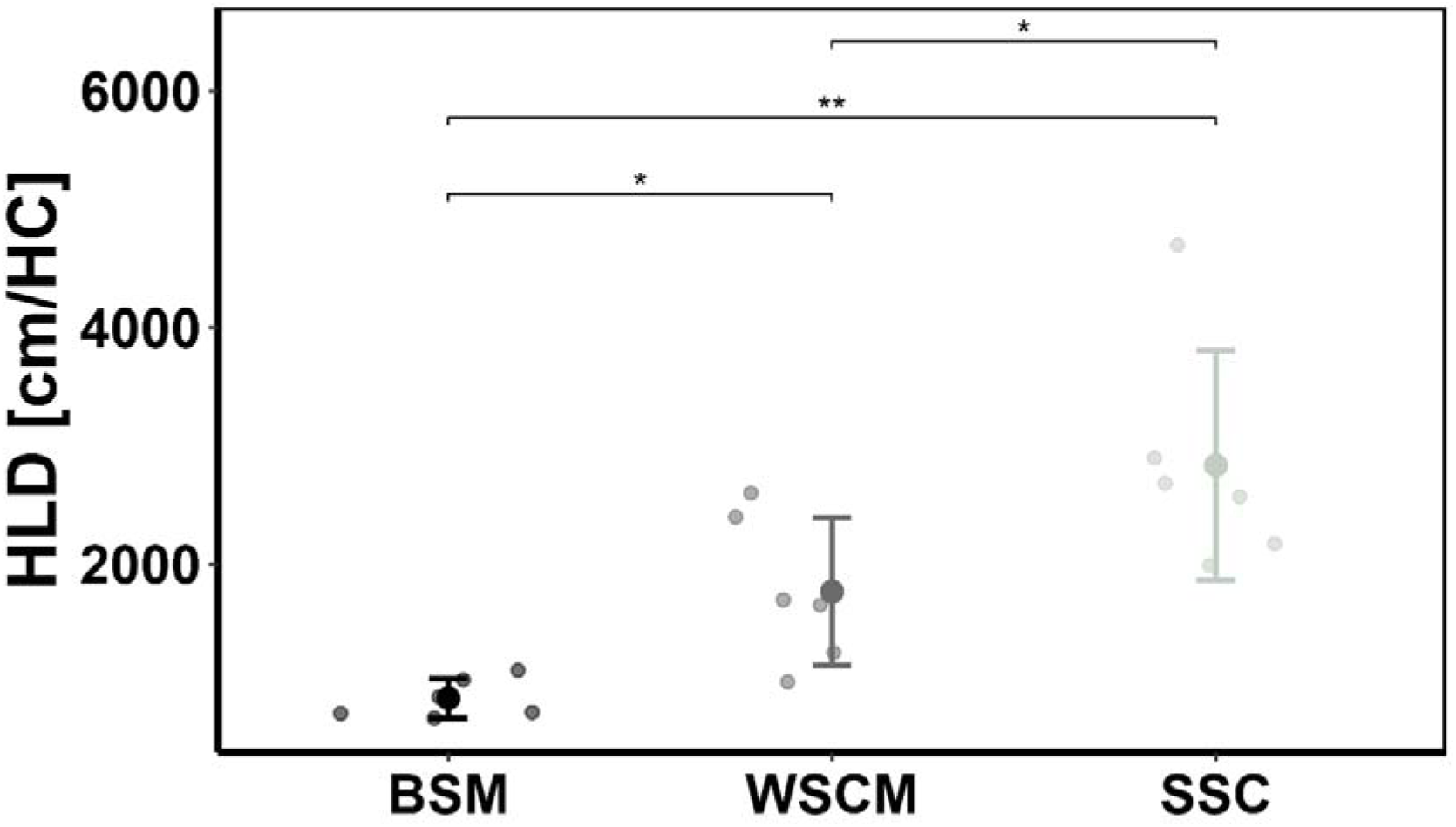
Comparison of fungal hyphal length density (HLD) in hyphal containers obtained by three different isolation methods. Soil from three hyphal containers (HC) was mixed and again separated into three equal parts. Fungal hyphae within these subsamples were extracted using either the modified aqueous blending and sieving method (BSM) (Jakobsen et al., 1992), the differential water/sucrose centrifugation method (WSCM) (Bingle and Paul, 1986), or our newly developed sieving and sucrose centrifugation (SSC) method. HLD was then measured using the visual gridline intersection (VGI) method. Statistical analysis was conducted using one-way ANOVA followed by Tukey’s HSD test (p<0.05, n=6). Significance codes: 0 ‘***’ 0.001 ‘**’ 0.01 ‘*’ 0.05 ‘.’ 0.1 ‘ns’ 1.

To assess the distribution of hyphae within the hyphal container, samples were collected from six distinct positions, spanning both horizontal and vertical orientations (Figure 3). Analysis using the SSC method revealed an even distribution of hyphae, which is essential when working with subsamples.

**Figure 3:**
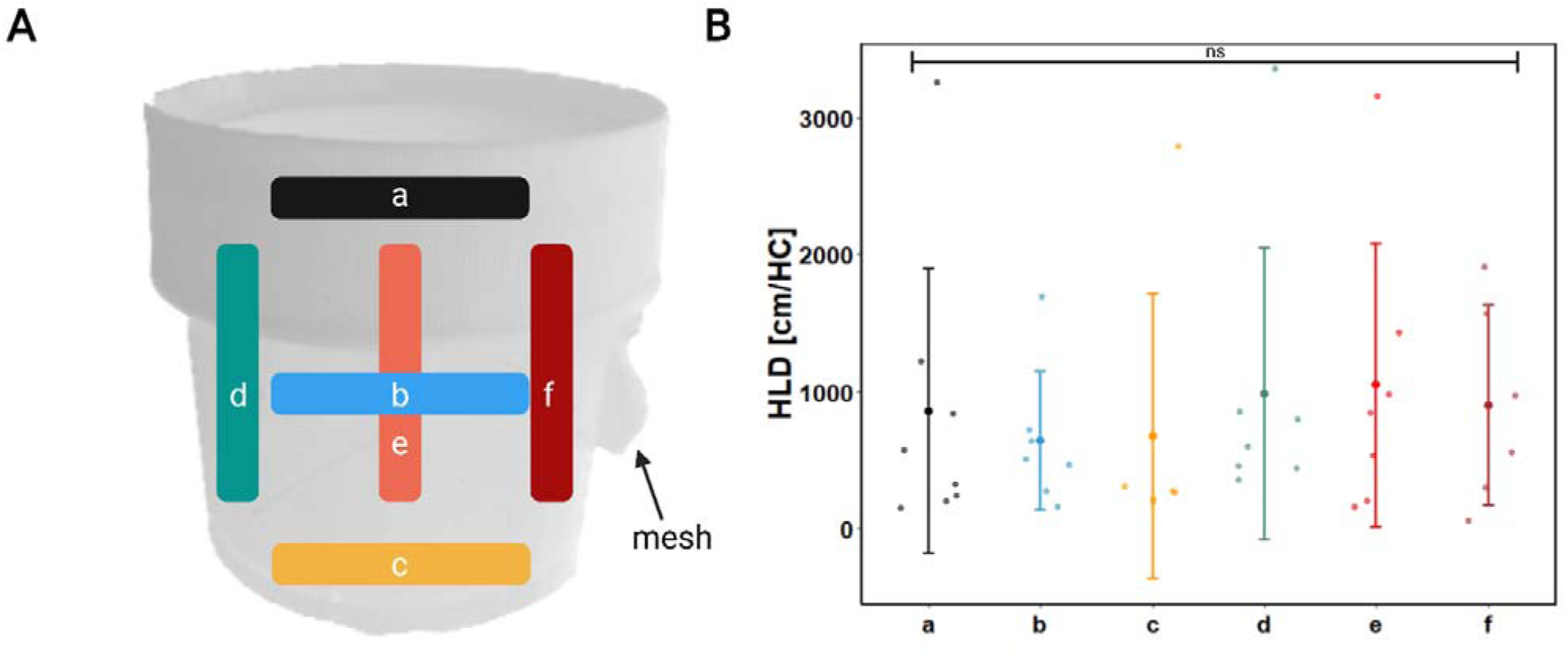
Hyphal length density is evenly distributed within the hyphal container. The hyphal containers harvested from field 2019 were divided into either three horizontal subsamples (top (a), middle (b), bottom (c)) or three vertical subsamples (left (d), middle (e), right (f)). Hyphal length density (HLD) within these subsamples was determined by the SCC and VGI methods. Different letters indicate significant differences between HLD in positions (one-way ANOVA followed by Tukey’s HSD test, p <0.05, n= 5-7). Significance codes: 0 ‘***’ 0.001 ‘**’ 0.01 ‘*’ 0.05 ‘.’ 0.1 ‘ns’ 1.

Next, the relationship between hyphal length density (HLD), measured after hyphal extraction with the SSC method, and fungal structures within maize wildtype roots was examined. Interestingly, HLD showed the strongest positive correlation (R = 0.5–0.8) with the combined presence of intraradical hyphae and arbuscules (H+A) within *Zea mays* root cells, followed by a slightly weaker correlation with total arbuscular mycorrhizal root colonization (including H+A and H+A+V), suggesting that extraradical hyphal growth is closely linked to stages of active arbuscular mycorrhizal symbiosis (Figure 4). However, at position 3 of the hyphal container - located in the topsoil and furthest from the root system center - this correlation was absent.

**Figure 4:**
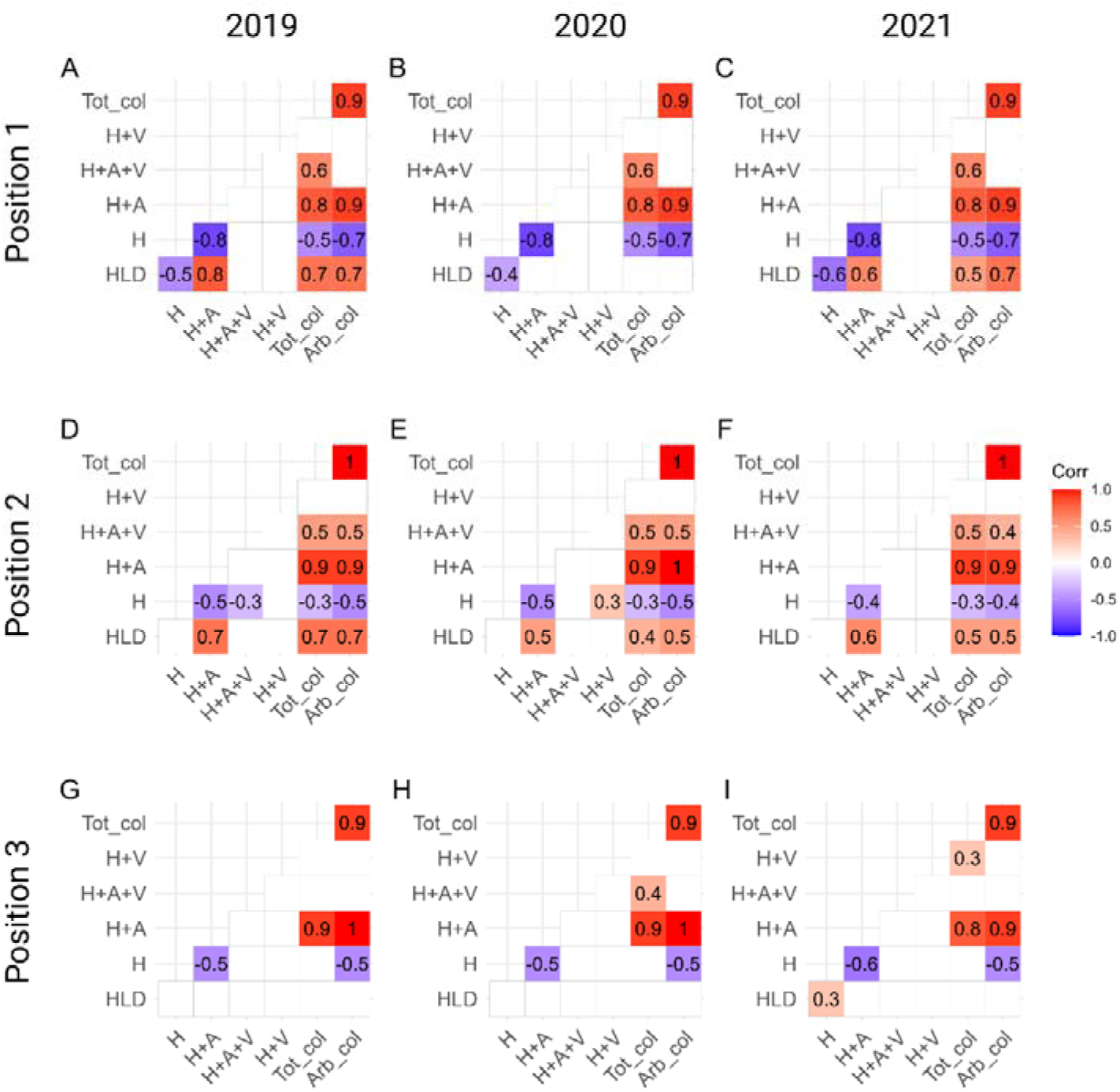
Hyphal length density correlates with the presence of hyphae and arbuscules (H+A). Hyphal containers were harvested from wildtype *Zea mays* plants grown on NK (-P), PK (-N), or NPK fields in 2019, 2020, and 2021, and hyphal length density within the hyphal containers was determined using the SCC and VGI methods. Fungal structures in roots were classified under the microscope into one of the following categories: hyphae alone (H), hyphae with vesicles (H+V), hyphae with arbuscules (H+A), hyphae, arbuscules, and vesicles (H+A+V), or non-colonized (no-myc). The percentage of fungal structures was correlated with HLD and with each other. Correlations are presented separately for each field year and for the containers at positions 1 to 3. Only significant correlations (p<0.05) are shown, with numbers indicating Spearman’s rank correlation coefficient. The colors represent positive and negative correlations, from red to blue. n=18.

Since AM fungi are not the dominant fungal subphylum in the hyphoplane samples and do not account for all extraradical hyphae extracted using the SSC method, these findings further suggest that active AM symbiosis influences the growth of a broader community of filamentous fungi (Figure 6C). This points to a potential, previously unrecognized feedback mechanism between intraradical colonization and extraradical hyphal proliferation. Moreover, the results indicate that the SSC method effectively estimates hyphal length.

Importantly, scanning electron microscopy, PCR-based detection, and DAPI staining confirmed that bacteria remained attached to fungal hyphae after extraction from field soil using the SSC method (Figure 5, Supplementary Figure S4). This demonstrated that the SSC method is well suited for studying bacterial hyphoplane communities in field experiments without the need for additives in natural substrates.

**Figure 5:**
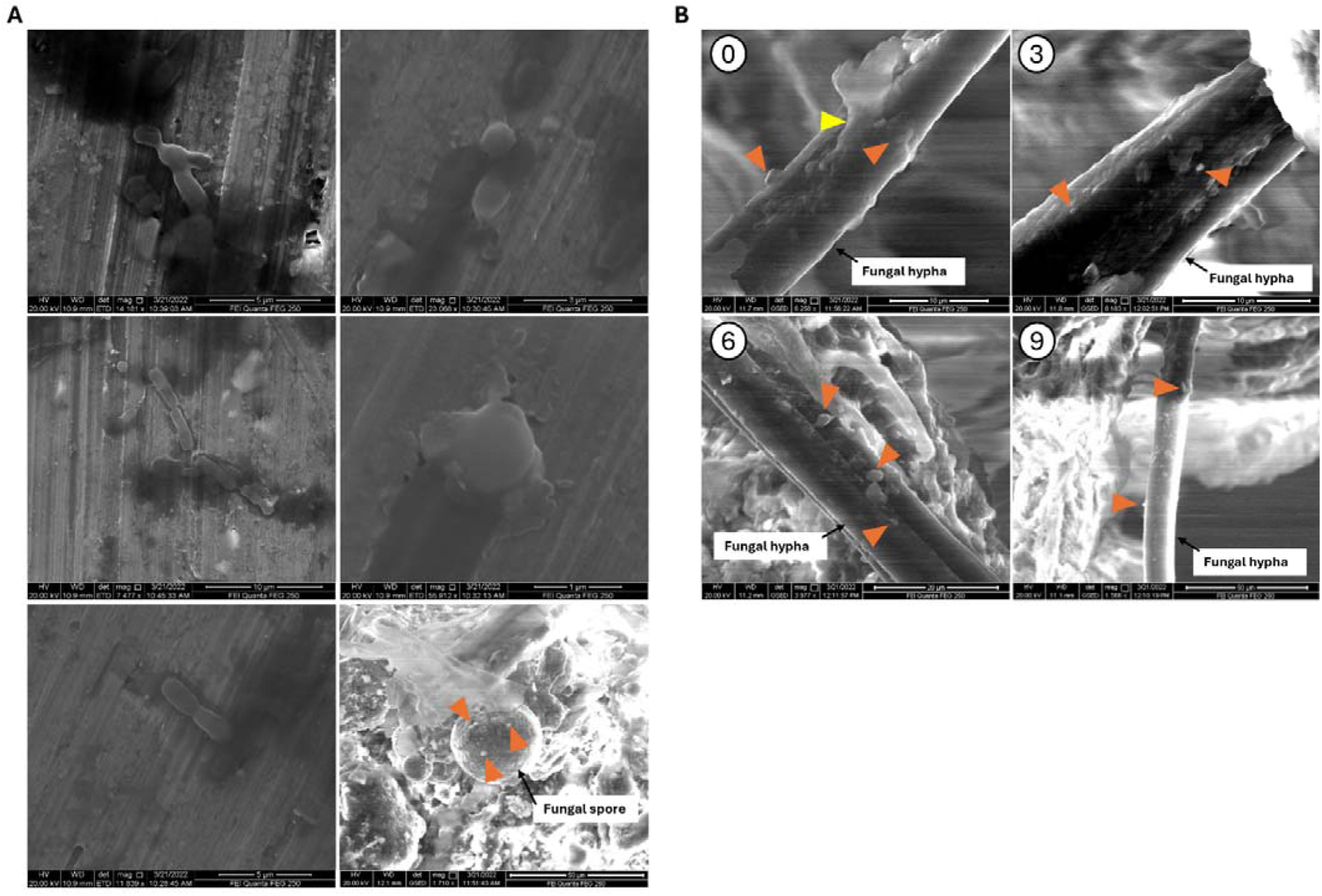
Scanning electron microscopy of microbial attachment to fungal hyphae after each critical step of the SSC method. (A) Scanning electron microscopy images taken of bacteria (control samples). (B) 0,3,6,9 indicate the steps of the SSC method before starting the method, after the second sieving step, after the last washing step and after lyophilization. Orange arrows indicate bacteria and yellow arrow shows bacterial biofilm.

Additionally, the SSC approach effectively enriched fungal hyphae from a highly diverse community sampled across three consecutive field experiments, with the phylum *Mucoromycota* and the genera *Minimedusa* and *Phialophora* dominating the hyphoplane (Figure 6C,D). However, the method does not selectively enrich AM fungal hyphae with attached microbes from field soil, as similarly sized hyphae from other fungal subphyla are also retained. As a result, identifying bacteria specifically associated with AM fungal hyphae and their role in nutrient cycling remains challenging. To exclude non-mycorrhizal fungal species, additional morphological examination of hyphae under a microscope would be required.

**Figure 6:**
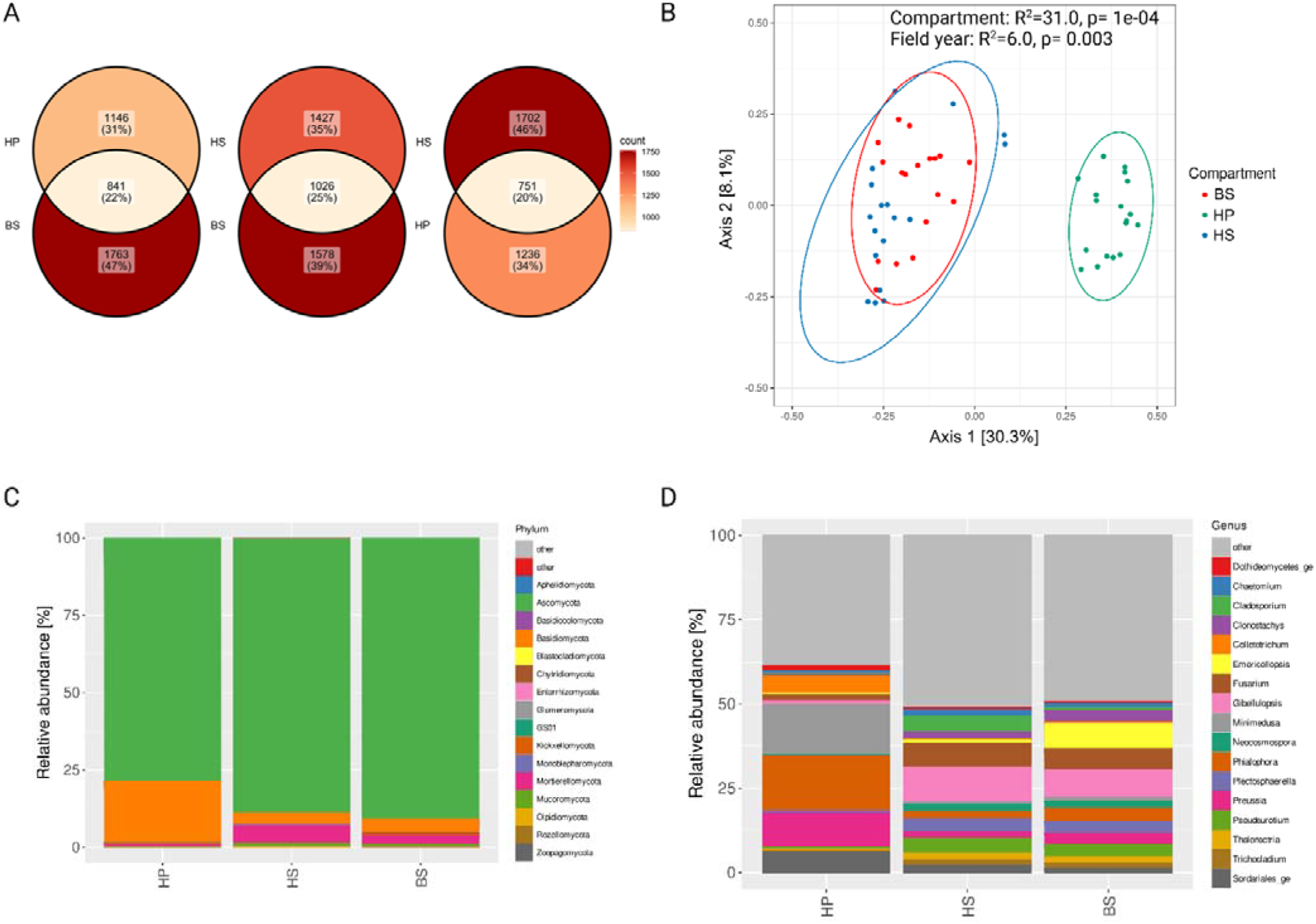
The SSC approach enriched fungal hyphae from a highly diverse fungal community. (A) Venn diagram depicting the number of ASVs and the percentage of shared fungal ASVs between compartments sampled from NK (-P) fields in 2019, 2020 and 2021. The color scale represents the number of ASV counts, ranging from bright orange (low number of ASVs) to dark red (high number of ASVs) (B) PCoA illustrating the Bray-Curtis dissimilarity of the fungal community compositions of hyphoplane (HP), hyphosphere (HS) at hyphal container position 1 and bulk soil from NK (-P) fields 2019, 2020 and 2021. Relative abundance of fungal (sub)phyla (C) and genera (D) in HP, HS and BS samples, respectively. n=18.

Our findings show that 22% of fungal ASVs were shared between the hyphoplane and bulk soil, suggesting that these fungi likely exhibit filamentous growth (Figure 6A). This also indicates that the SSC method successfully enriched 22% of the total soil fungal diversity by specifically targeting filamentous fungi. For bacteria, 55% of the taxa detected in bulk soil were also found in the hyphoplane, while this proportion was 4% higher in the hyphosphere (Figure 7A). The overlap of ASVs between the hyphosphere and hyphoplane was 1–2% lower than that between the hyphoplane and bulk soil.

**Figure 7:**
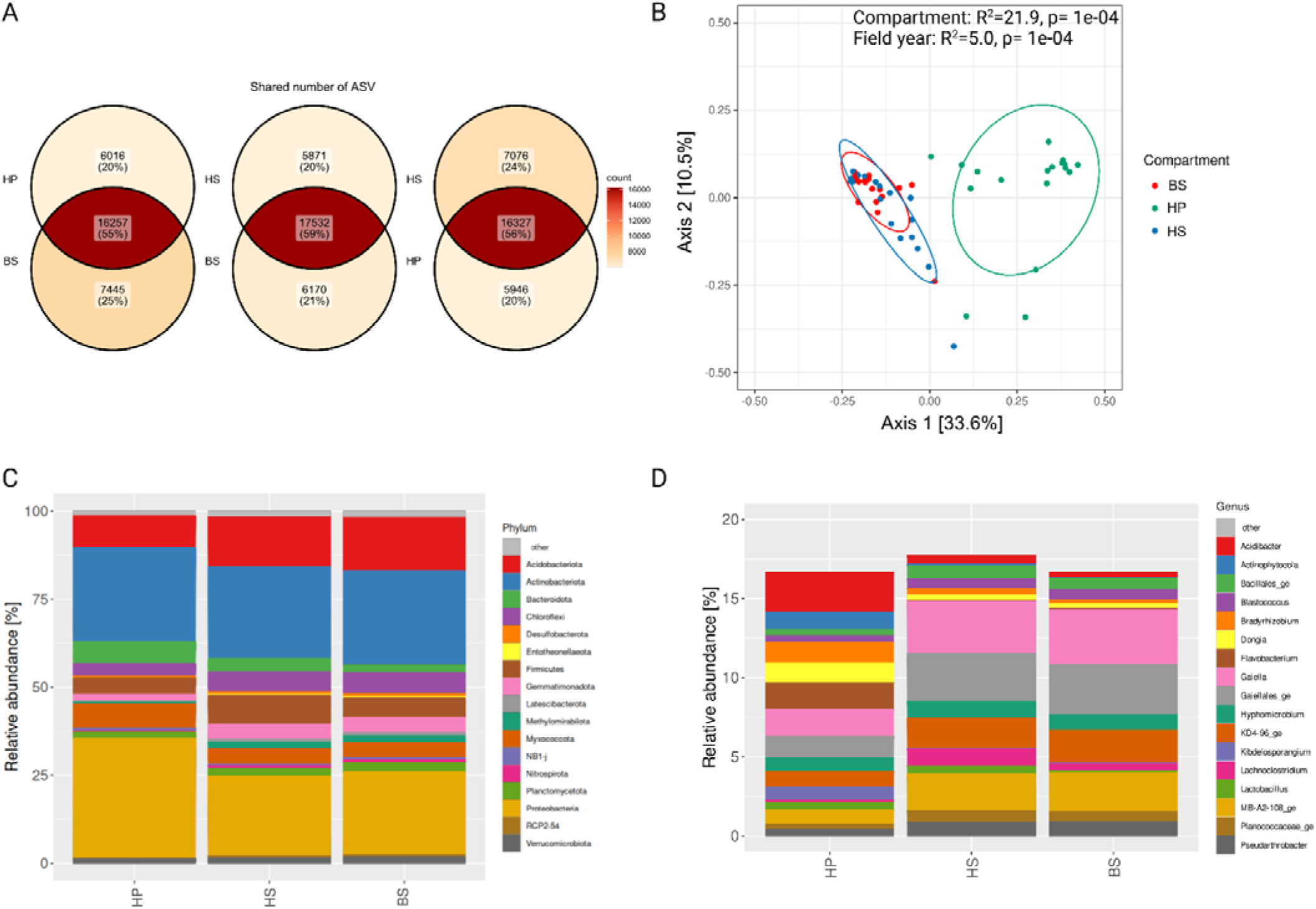
Bacterial community composition differs between hyphoplane and soil compartments. (A) Venn diagram depicting the number of ASVs and the percentage of shared bacterial ASVs between compartments sampled from NK (-P) fields in 2019, 2020 and 2021. The color scale represents the number of ASV counts, ranging from bright orange (low number of ASVs) to dark red (high number of ASVs). (B) PCoA illustrates the Bray-Curtis dissimilarity of the bacterial community compositions of hyphoplane (HP, Green), hyphosphere (HS, Blue) at hyphal container position 1 and bulk soil (BS, Red) from NK (-P) fields 2019, 2020 and 2021. Relative abundance of bacterial phyla (C) and genera (D) in HP, HS and BS samples, respectively. n=18.

To further assess differences in fungal and bacterial community composition across compartments, beta diversity was analyzed. Principal Coordinates Analysis (PCoA) revealed that fungal and bacterial communities in the hyphoplane were clearly distinct from those in bulk soil and the hyphosphere, whereas bulk soil and hyphosphere communities exhibited substantial overlap (Figure 6B, 7B). This suggests that conditions within the container closely resemble those of natural soil environments. PERMANOVA confirmed that beta diversity was primarily affected by compartment type (31.0% and 21.9% of the variance for fungi and bacteria, respectively) and less by the year in which the experiment was conducted (6.0% and 5.0%), suggesting consistent patterns across field years.

The bacterial hyphoplane community was dominated by the phyla Proteobacteria and Actinobacteriota (Figure 7C), aligning with findings from previous hyphoplane studies (Emmet et al., 2021; Wang et al., 2022; Zhang et al., 2024). The genera *Acidibacter* and *Flavobacterium* were the most relatively abundant in the hyphoplane (Figure 7D). However, detailed analyses of microbiome × environment × genotype interactions will be presented elsewhere.

It is important to note that these communities may also include microbes residing within the hyphal endosphere, although they are likely a minority, as only a few specific taxa have been identified as hyphal endosymbionts. For example, *Candidatus Glomeribacter gigasporarum*, a rod-shaped, Gram-negative beta-proteobacterium, has been consistently associated with Glomeromycotina (Bonfante et al., 1994; Salvioli et al., 2016). Additionally, Glomeromycotina host round-shaped bacteria resembling Gram-positive bacteria, known as Mollicutes-related endobacteria (Naumann et al., 2010). Future studies will enable a clearer experimental distinction between bacteria inhabiting the hyphal endosphere and those colonizing the hyphoplane, providing deeper insights into their ecological roles.

To ensure high DNA quality and minimize microbial community changes during sample processing, we performed the SSC method at a maximum of 4°C. Whether this method also ensures sufficient RNA quality remains to be determined.

In conclusion, this study presents a rapid and efficient method that, compared to previously described approaches, is semi-automated and enables both the extraction of hyphae and the identification of hyphoplane bacteria from natural substrates. By streamlining these processes, this methodology enables the identification of hyphoplane bacteria in large-scale field studies and provides a valuable tool for investigating the effects of soil management and plant genotype on hyphoplane communities.

## Abbreviations

AMF: arbuscular mycorrhizal fungi
ERH: extraradical hyphae
HC: hyphal compartment
NGS: next-generation sequencing
Pi: inorganic orthophosphate
SSC: sieving and sucrose centrifugation
VGI: visual gridline intersection

## 5 Acknowledgements

We thank Heinz Georg Moheim and Dr. Frank Nitsche for assistance with the set-up of the water pump and scanning electron microscopy, respectively.

## 6 Author contributions

IM, MB: conceptualization; IM: methodology; IM: writing - original draft; IM and MB: writing - review & editing; IM, MB: visualization

## 9 Data Availability

All primary data to support the findings of this study will be made available on request.

---

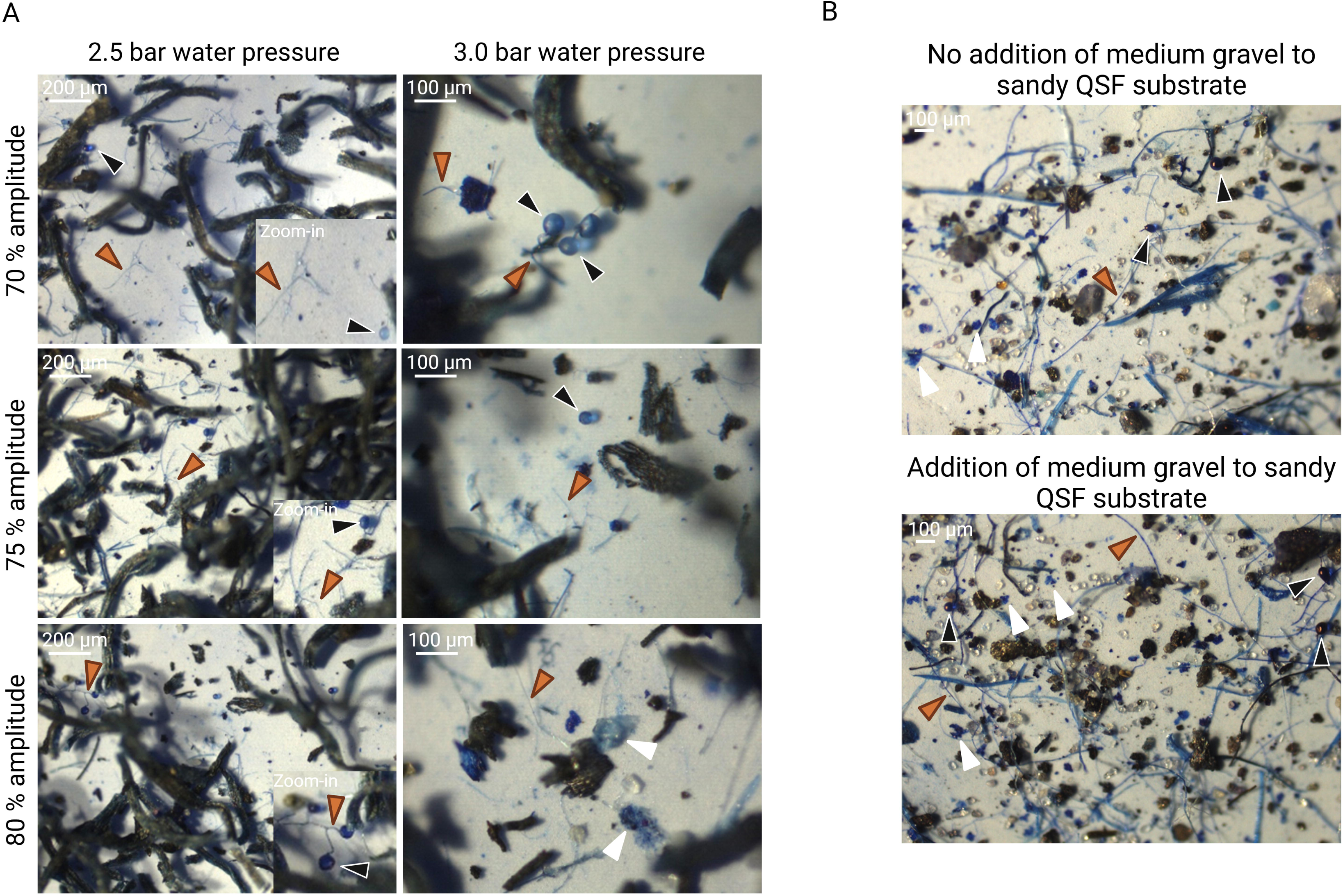

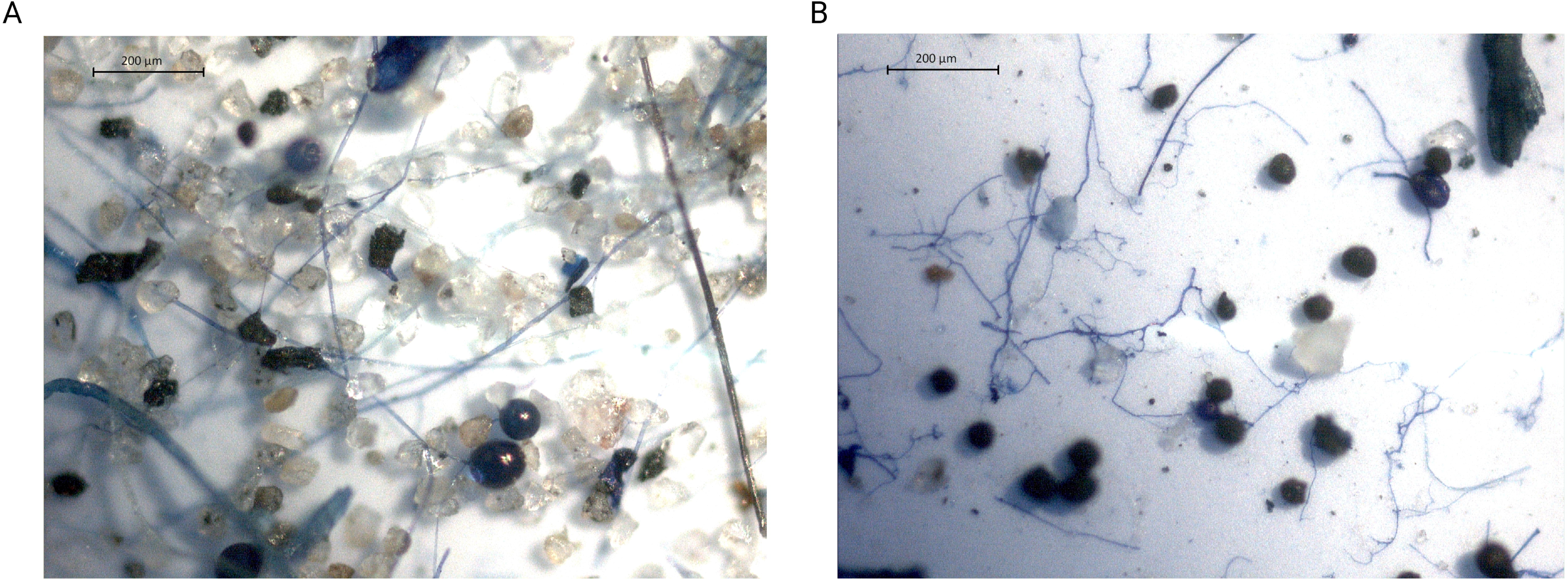

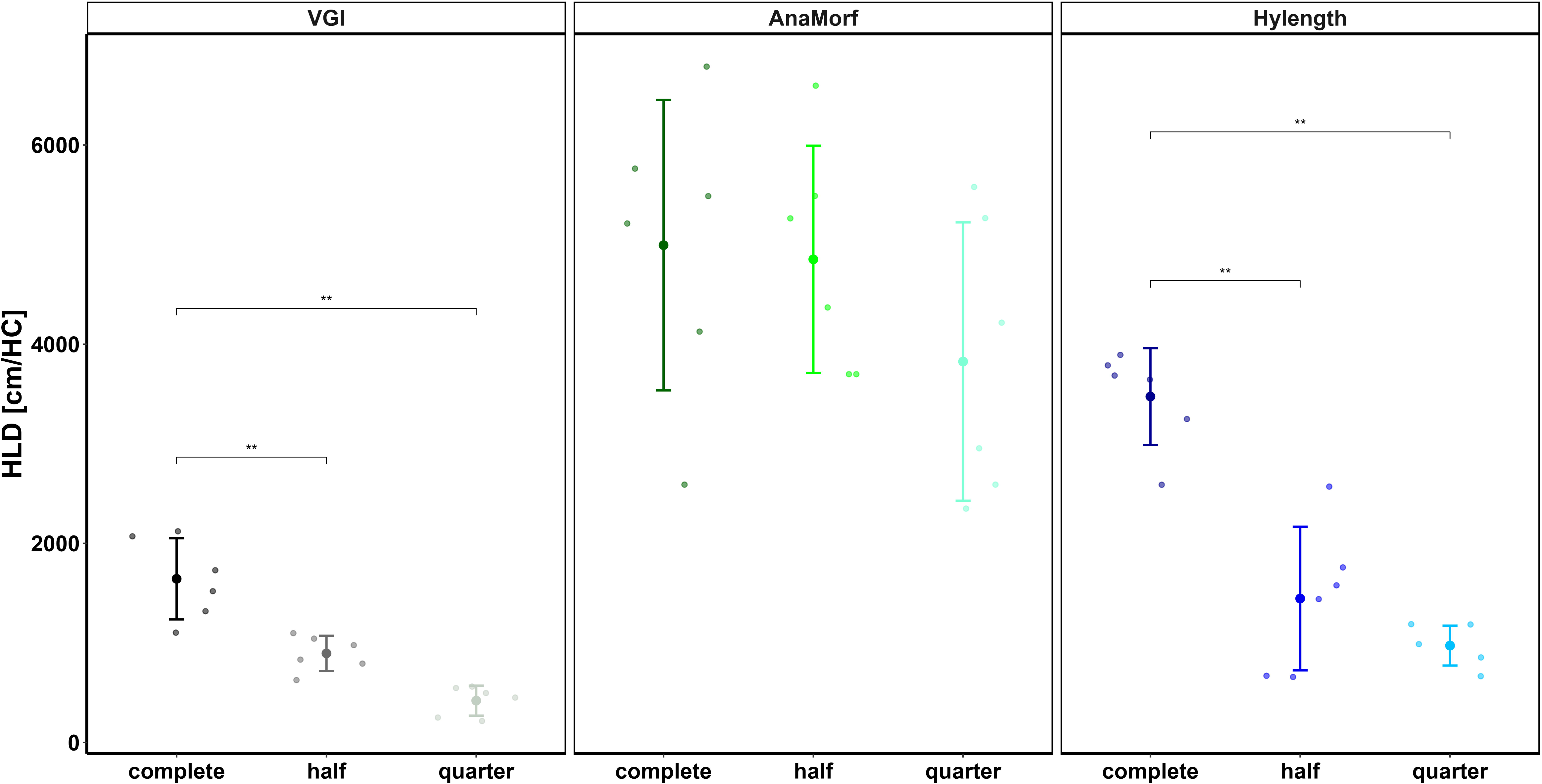

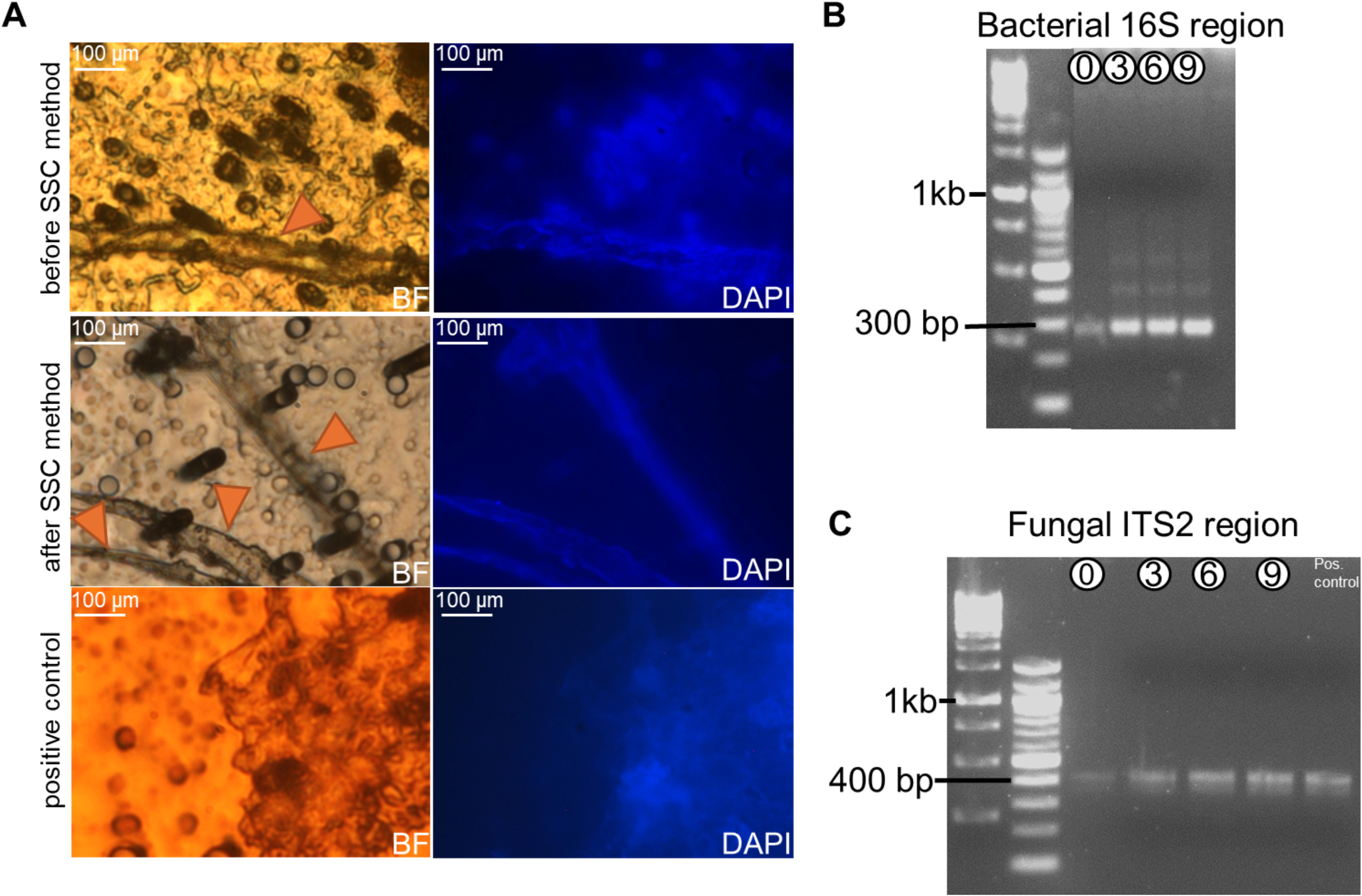

